# Noise propagation in gene expression in the presence of decoys

**DOI:** 10.1101/2020.04.01.020032

**Authors:** Supravat Dey, Abhyudai Singh

**Affiliations:** Department of Electrical and Computer Engineering, University of Delaware, Newark, DE 19716, USA; Department of Electrical and Computer Engineering, Department of Biomedical Engineering and Department of Mathematical Sciences, University of Delaware, Newark, DE 19716, USA

## Abstract

Genetically-identical cells can show remarkable intercellular variability in the level of a given protein which is commonly known as the gene expression noise. Besides intrinsic fluctuations that arise from the inherent stochasticity of the biochemical processes, a significant source of expression noise is extrinsic. Such extrinsic noise in gene expression arises from cell-to-cell differences in expression machinery, transcription factors, cell size, and cell cycle stage. Here, we consider the synthesis of a transcription factor (TF) whose production is impacted by a dynamic extrinsic disturbance, and systematically investigate the regulation of expression noise by decoy sites that can sequester the protein. Our analysis shows that increasing decoy numbers reduce noise in the level of the free (unbound) TF with noise levels approaching the Poisson limit for large number of decoys. Interestingly, the suppression of expression noise compared to no-decoy levels is maximized at intermediate disturbance timescales. Finally, we quantify the noise propagation from the TF to a downstream target protein and find counterintuitive behaviors. More specifically, for nonlinear dose responses of target-protein activation, the noise in the target protein can increase with the inclusion of decoys, and this phenomenon is explained by smaller but more prolonged fluctuations in the TF level. In summary, our results illustrates the nontrivial effects of high-affinity decoys in shaping the stochastic dynamics of gene expression to alter cell fate and phenotype at the single-cell level.

## I. INTRODUCTION

Genetically identical cells can show remarkable variability in the level of a gene product (mRNA/protein) which is commonly known as the gene expression noise [1]–[9]. Depending on the situation, the gene-expression noise can be detrimental or beneficial [10]–[14]. For example, large noise can cause defects in developing embryos [15]. Gene expression noise can drives different cell-fates of genetically identically cells [12]. Importantly, it can enhance phenotypic diversity, crucial for the survival of an organism in a population under fluctuating environments [13], [14].

There are two components of the cell-to-cell variability in gene expression — intrinsic and extrinsic [2], [3], [6], [7], [15]. The intrinsic noise arises from the inherent stochasticity of biochemical reactions associated with mRNA/protein productions and degradations involving few molecules. There are several cell-specific factors such as, cell cycle stage, cell size, abundance of expression machinery and global factors that are the extrinsic sources of expression noise [16]– [20]. The extrinsic and intrinsic noises can be quantified by experiments with two-color reporter genes that are regulated by identical promoters [2]. While correlated fluctuations represent the extrinsic noise, the uncorrelated fluctuations is a measure of intrinsic noise [2].

Here, we investigate the role of genomic decoy binding on the gene expression noise in the presence of a dynamic external disturbance. The genomic decoys are the numerous high-affinity binding sites where a transcription factors (TFs) bind without any direct functional consequences [21]. In functional binding, transcription factors bind to the specific site of a gene and *directly* regulate the expression by activating or inhibiting the transcription process. The binding of transcription factors to the decoy sites, on the other hand, alters the abundance of the transcription factor and *indirectly* regulates the expression of target genes. The role of decoys in the stochastic gene expression were studied experimentally using synthetic circuits in *Saccharomyces cerevisiae* and in various non-oscillatory circuits [22]–[27] and oscillatory circuits [28], [29] theoretically. However, the effect of decoys in the presence of external disturbance has not been addressed.

To investigate the role of decoys on the gene expression noise in the presence of a dynamic external perturbation, we formulate a simple stochastic model for the gene expression, schematically shown in Fig. 1. The transcription factor, whose production rate is subject to an external perturbation, reversibly binds to unoccupied decoy sites. Assuming small fluctuations in molecular copy numbers, we linearize associated binding terms around the means and solve the first and second-order moment dynamics at the steady-state. We quantify the noise in free TFs using the squared coefficient of variation and obtain analytical formulas for noise levels in the fast binding/unbinding limit. Our results show that the addition of decoy sites reduces the extrinsic noise component of the free TF level. Finally, using stochastic simulations, we investigate the noise for a downstream protein. Interestingly, we observe that decoys can enhance or buffer noise in the target protein depending on the nature of the dose response of the target protein activation.

**Fig. 1.**
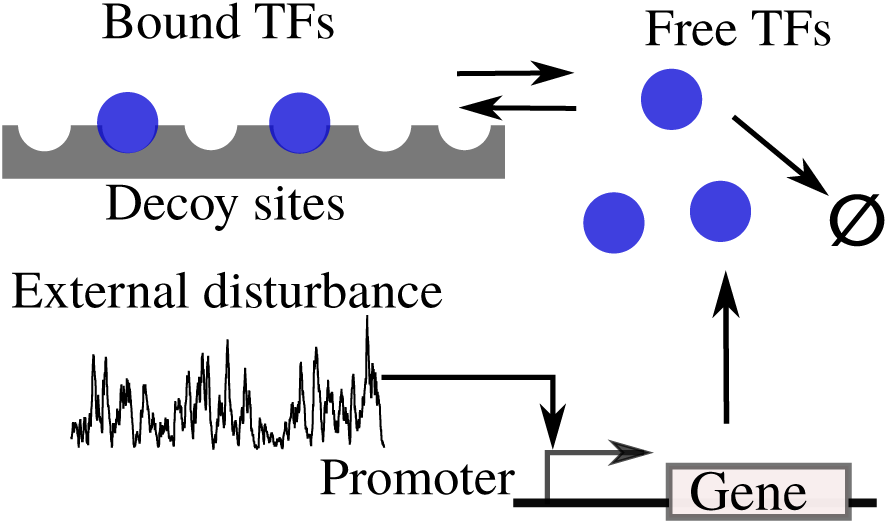
Model schematic of TF expression impacted by upstream extrinsic noise. The synthesis of a transcription factor (TF) from a gene is modeled as a simple birth-death process. An upstream disturbance affects the stochastic dynamics of TF copy numbers by making the synthesis rate itself a random process. There are *N* decoy binding sites in the genome that can sequester the TF. A free TF molecule reversibly binds to an unoccupied decoy site with relatively fast binding/unbinding kinetics compared to the timescales of disturbance and TF turnover.

### Symbols and Notation

At a given time *t*, the number of molecules for the species associated with the external disturbance is denoted by *x*(*t*), the molecular number of free and bound TFs by *y*_*f*_ (*t*) and *y*_*b*_(*t*), and the number of target protein by *z*(*t*). The molecular count of a species takes a random non-negative integer value i.e., *x*(*t*), *y*_*f*_ (*t*), *y*_*b*_(*t*), *z*(*t*) ∈ {0, 1, 2, 3, …}. For a stochastic process, we use angular brackets ⟨.⟩ and 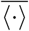 to denote the transient and steady state expected values respectively. The total noise in the molecular counts of a free TF is quantified by the square coefficient variation at the steady-state

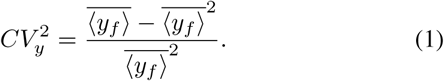

Similarly, we quantify noise in external disturbance and target protein count using 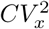 and 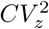, respectively.

## II. Modeling Gene expression in the presence of extrinsic disturbances and decoy sites

We formulate a simple stochastic model schematically shown in Fig. 1, where the synthesis of TF is subject to an extrinsic disturbance. This disturbance biologically corresponds to fluctuations in the abundance of enzymes, expression machinery, or other global factors connected to cell size/cell-cycle stage. As discussed in further detail below, we phenomenologically model this disturbance as a bursty birth-death process.

### A. Modeling extrinsic disturbance as a bursty process

Increasing evidence shows that expression of gene products in single cells occurs in intermittent bursts [30]–[37], and bursty birth-death processes have been commonly used to capture fluctuations in the levels of these gene products [38]–[43]. Borrowing this framework, we model the extrinsic disturbance as a chemical species that is produces in bursts as per a Poisson process with rate *k*_*x*_, and the species is subject to degradation at a constant rate. Let *x*(*t*) denote the level of the disturbance at time *t*. Then, whenever a burst event occurs the level is reset by

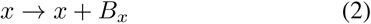

where the burst size *B*_*x*_ is an independent and identically distributed random variable with *Prob*(*B*_*x*_ = *i*) = *α*(*i*), *i ∈* {1, 2, …}. Similarly, when a degradation event occurs the levels are reset as

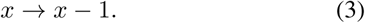

The probabilities of these rests occurring in the next infinitesimal time interval (*t, t* + *dt*) are

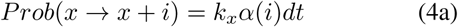

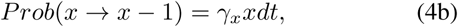

respectively, where *γ*_*x*_ is the decay rate. For this system the mean and noise in molecular copy number *x*(*t*) at steady state is given by

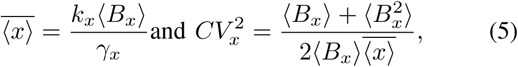

respectively, where ⟨*B*_*x*_⟩ is the average burst size [44], [45]. If ⟨*B*_*x*_⟩ follows a geometric distribution with mean ⟨*B*_*x*_⟩, then the second equation in (5) reduces to

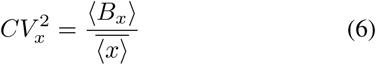

and the magnitude of disturbance fluctuations increases with increasing ⟨*B*_*x*_⟩. The speed of the fluctuations is given by the auto-correlation function

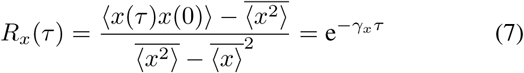

[43], [44]. Thus, the degradation rate *γ*_*x*_ is a measure of the speed or timescale of disturbance fluctuations. In other words, the fluctuations become fast as *γ*_*x*_ → *∞*, and become slow as *γ*_*x*_ → 0. Next, we describe how this disturbance impacts the synthesis of a TF.

### B. Effect of extrinsic noise in the absence of decoy

First, we discuss the effect of extrinsic noise on the stochastic synthesis of the TF. The dynamics of TF is assumed to follow a simple birth-death process with production rate that linearly depends on the disturbance, and a constant decay rate *γ*_*y*_. Analogous to (4), but assuming no bursting, the probability of resets in the levels of the TF occurring in the next infinitesimal time interval (*t, t* + *dt*) are

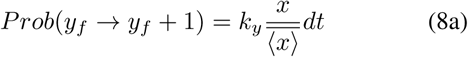

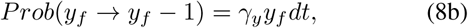

where *y*_*f*_ (*t*) denotes the level of the TF at time *t*. To analyze the coupled random processes (8) and (4) we use the framework of moment dynamics for discrete-state continuous-time Markov processes as described in [46]. More specifically, the time evolution of any arbitrary function 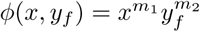 for *m*_1_, *m*_2_ *∈* {0, 1, 2, …} is given by,

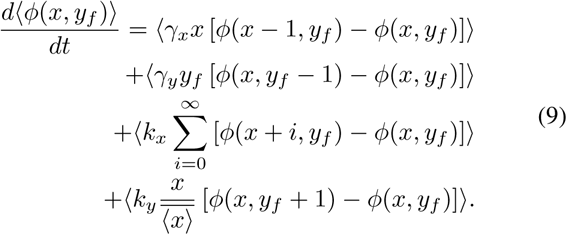

We use this above extended generator to obtain the time evolution of all the first and second-order moments of *x*(*t*) and *y*_*f*_ (*t*). Solving these moment dynamics at steady-state reveals the following noise in the level of the TF (as quantified by its coefficient of variation squared)

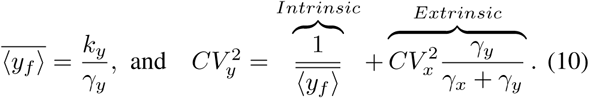

The first part of 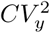 is the intrinsic noise due to the Poissonian birth-death process, and the second part is the additional contribution due to the extrinsic noise. For a very fast external perturbation, *γ*_*x*_ → *∞*, the noise is purely intrinsic. For a very slow disturbance *γ*_*x*_ → 0, 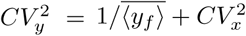.

### C. Effect of extrinsic noise in the presence of decoy

Next, we expand the model to include *N* decoy binding sites. The free TF reversibly binds to unoccupied decoy sites with binding and unbinding rates *k*_*b*_ and *k*_*u*_, respectively. While a free TF molecule degrades at constant rate *γ*_*y*_, we assume that TFs bound to decoys are protected from degradation [23], [25], [27].

Let random processes *y*_*f*_ (*t*) and *y*_*b*_(*t*) denote the level of free and bound TF. Then, the probability of resets in *y*_*f*_ (*t*) and *y*_*b*_(*t*) corresponding to different events occurring in (*t, t* + *dt*) are as follows

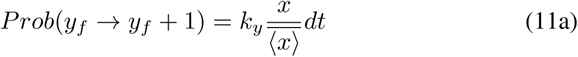

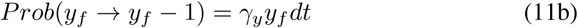

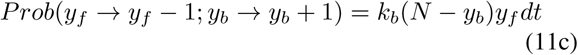

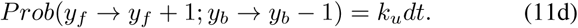

Here the first two resets represent the synthesis and degradation of a free TF molecule, and the last two resets represent binding/unbinding to decoys. Note that *N* − *y*_*b*_ is the number of unoccupied decoys resulting in the *nonlinear* binding rate is *k*_*b*_(*N* − *y*_*b*_)*y*_*f*_. Combing (11) with (4) yields the following extended generator for any arbitrary function *φ*(*x*_*f*_, *y*_*f*_, *y*_*b*_):

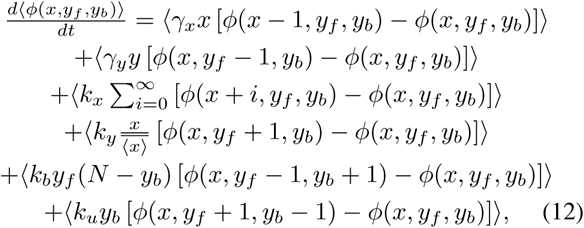

and time evolution of moments is obtained by setting 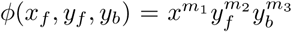 for *m*_1_, *m*_2_, *m*_3_ *∈*{0, 1, 2, …} [46]. It turns out that the nonlinear binding rate results in the problem of unclosed dynamics – time evolution of lower order moments depends on higher order moments, and typically moment closure schemes are exploited to compute moments. Here we use the well-known linear noise approximation [48]–[51] that under the assumption of small fluctuations in *y*_*f*_ and *y*_*b*_ around their respective mean values ⟨*y*_*f*_ ⟩ and ⟨*y*_*b*_⟩, linearizes the nonlinearity as

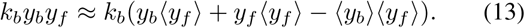

Using these linearized rates in place of the nonlinearity *k*_*b*_*y*_*b*_*y*_*f*_ in (12), one can again derive the time evolution of the moments and solve them to get approximate analytical formulas for the TF noise levels. Given the space constraints, we omit the proof and only present the main results.

Solving the first order moment dynamics at the steady-state, we obtain the expressions for the mean free TF and bound TF counts,

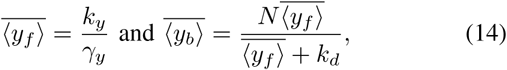

where, *k*_*d*_ = *k*_*u*_*/k*_*b*_ is the dissociation constant. Please note that 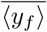 is independent of the decoy sites as has been shown before in the absence of disturbance [25]. Solving the second order moment dynamics, we obtain the following formula for the noise in the free TF in the limit of fast binding/unbinding (*k*_*b*_ → *∞, k*_*u*_ → *∞* and *k*_*d*_ = *k*_*u*_*/k*_*b*_ being finite),

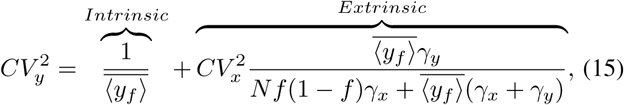

where 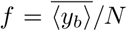 is the fraction of bound TFs. Note that decoy binding directly affect the extrinsic noise part. The noise decreases monotonically to the Poisson limit 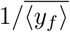 as *N* → *∞*. Fig. 2(A) shows how the noise decays with *N* for various values of the speed of the extrinsic (disturbance) fluctuations *γ*_*x*_. For a given *N*, 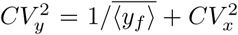 for *γ*_*x*_ *<< γ*_*y*_ and 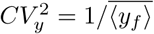 for *γ*_*x*_ *<< γ*_*y*_ (see Fig. 2(B)). If we plot the noise in free TF normalized with that of *N* = 0 as a function of *γ*_*x*_, the noise show a minima for a specific value of *γ*_*x*_ (see Fig. 2(C)). In essence, the noise suppression ability of decoys is highest at intermediate timescales of disturbance fluctuations.

**Fig. 2.**
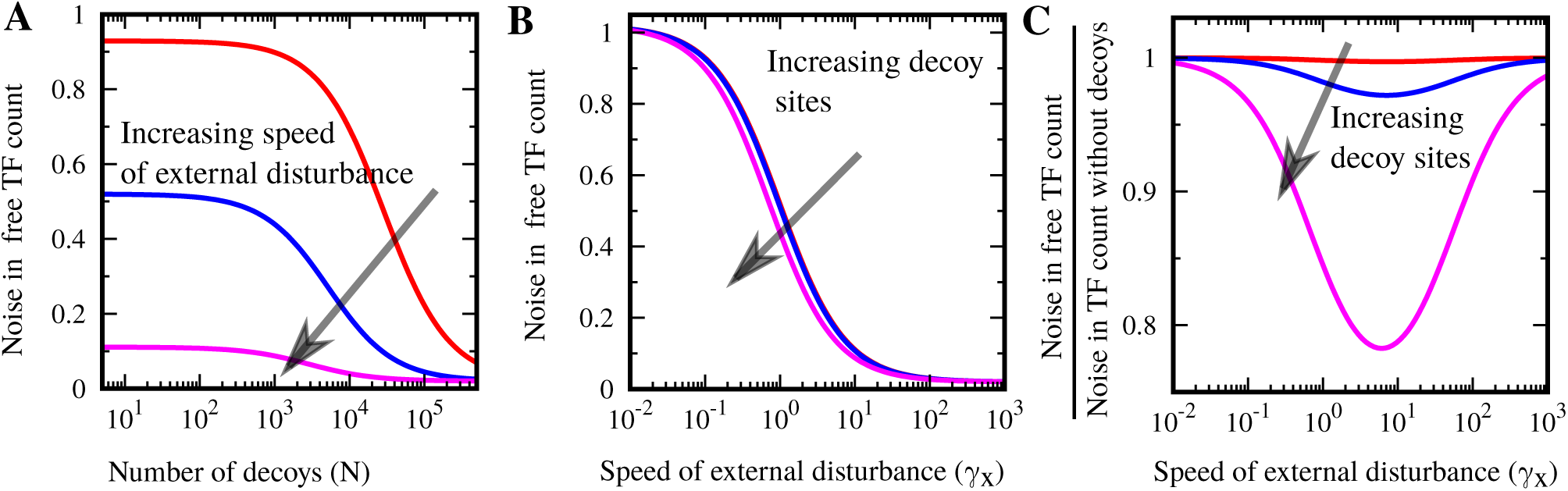
Decoy binding buffers gene expression noise due to extrinsic disturbance. (A) The noise in the free TF counts is plotted against *N* for different values of speed of the external disturbance (*γ*_*x*_ = 0.1, 1, and 10). The noise at *N* = 0 is higher for a slower external disturbance. For large *N*, the noise approaches the Poisson limit. (B) The noise in the free TF counts is plotted against *γ*_*x*_ for *N* = 10, 100, and 1000. The noise approaches to the Poisson limit for *γ*_*x*_ *>> γ*_*y*_. (C) An optimal noise behavior is observed when the noise normalized by the noise level when *N* = 0. The noise buffering becomes minimum for a particular value of *γ*_*x*_. This minimum value of the noise decreases with increasing *N*. Parameter used: ⟨*B*_*x*_⟩ = 50, 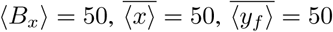, *γ*_*y*_ = 1, and *k*_*d*_ = 1.

## III. Noise in the Target protein

Having quantified the effect of decoys on TF expression noise, we next study noise propagation from the TF to a downstream target gene. Towards that end, free TF molecules activate the production of a target protein with synthesis rate *k*_*z*_*g*(*y*_*f*_) (Fig. 3(A)). We study two specific cases: (I) Linear activation, *g*(*y*_*f*_) = *k*_*z*_*y*_*f*_ and (II) Step-like activation (activation only occurs when 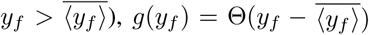, where Θ is the Heaviside step-function, i.e., Θ(*x*) = 0 when *x <* 0 and Θ(*x*) = 1 when *x >* 0. As in the case of the TF, assuming a non-bursty production of the target protein results in the following probability of resets

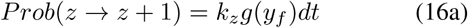

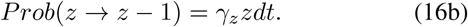

for the birth and death of the target protein, where *z*(*t*) denotes the level of the target protein, and *γ*_*z*_ is the target protein’s degradation rate. An important point to mention is that only the TF binds to decoys. A key focus here is to quantify the extent of noise propagation from the TF to the target protein as a function of decoy numbers for different dose responses *g*(*y*_*f*_).

**Fig. 3.**
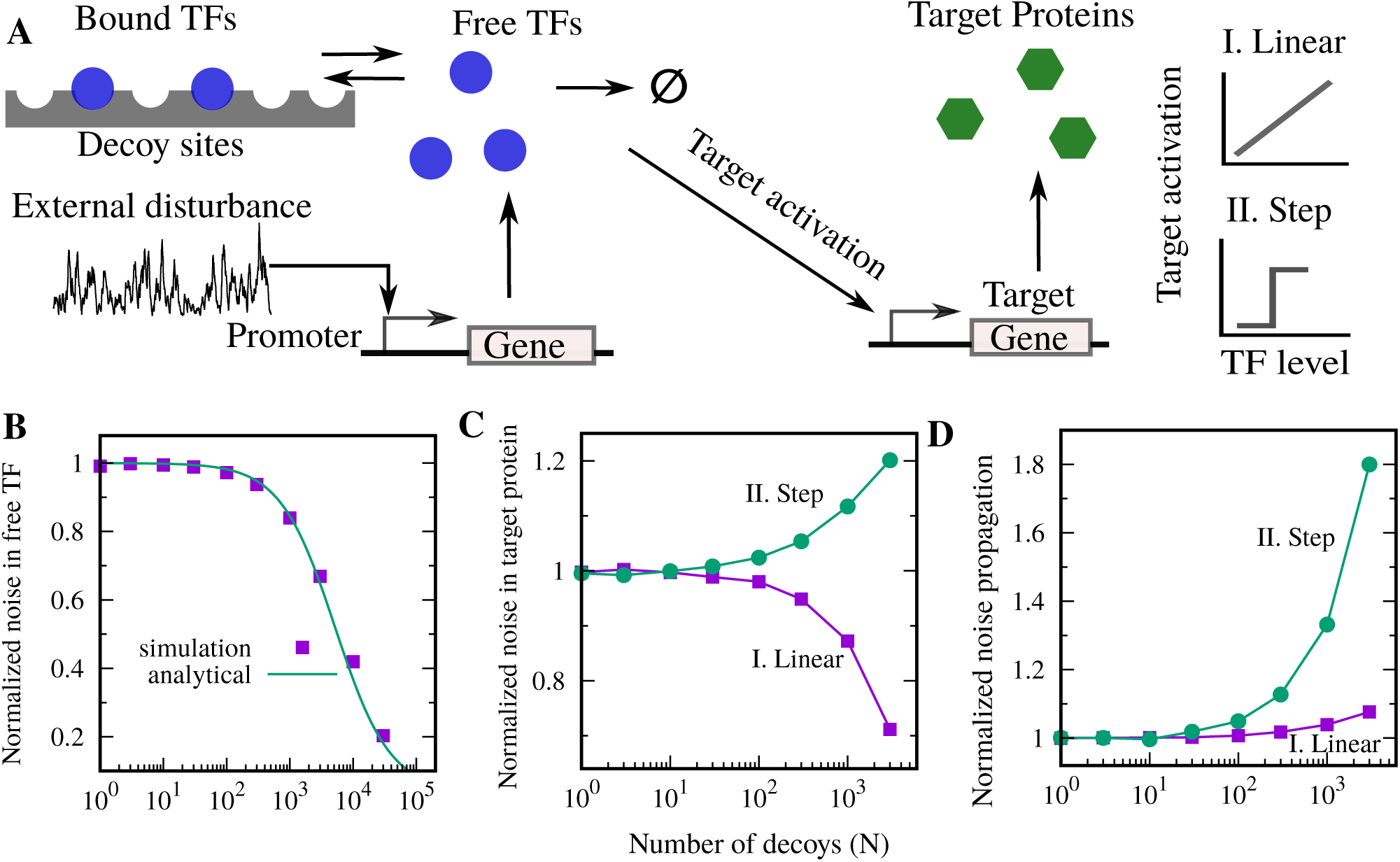
Target protein noise from stochastic simulations shows a distinct behavior depending on linear and step-like regulation. (A) A schematic diagram of the model for studying noise in the downstream gene expression. Free TFs activate the expression of the target gene: production rate is *k*_*z*_*g*(*y*_*f*_). We consider two cases: (I) Linear target activation: *g*(*y*_*f*_) = *y*_*f*_, and (II) Step-function target activation: 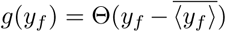, where Θ is the Heaviside step-function, i.e., Θ(*x*) = 0 when *x <* 0 and Θ(*x*) = 1 when *x >* 0. (B) The results of the noise in free TF counts obtained from running a large number of Monte Carlo simulations using the Stochastic Simulation Algorithm [47] and analytical formula (15) are in agreement. The noise in the target protein (normalized by 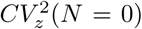) plotted against *N*. While for the linear regulation, the noise in the target protein decreases, for the step-like regulation noise in the target protein increases as a function of *N*. Noise propagation defined as the ratio of free TF noise level to the target protein noise level increases as function of *N*. However, the noise propagation is higher for the step-like dose response. For this plot parameters taken as ⟨*B*_*x*_⟩ = 50, *k*_*x*_ = 1, *k*_*y*_ = 50, *γ*_*x*_ = *γ*_*y*_ = *γ*_*z*_ = 1, *k*_*d*_ = 1, and *k*_*z*_ = 1.

We study this system of four nonlinearly coupled random processes *x*(*t*), *y*_*f*_ (*t*), *y*_*b*_(*t*) and *z*(*t*) by running a large number of Monte Carlo simulations using the Stochastic Simulation Algorithm [47]. Quantification of steady-state level noise levels for different species are shown Fig. 3. First, we show that the analytical result of noise in the free TF (15) as obtained using the linear noise approximation matches nicely with the simulations (Fig. 3(B)). The noise in the target protein 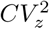 shows a counterintuitive behavior. Given that decoys buffers noise in free TF, one might expect the same behavior in the target protein noise. However, we see two opposite role of decoys depending on the profiles of the activation. For a linear dose response, decoys buffer noise in the target protein, but enhance noise for the step-like dose response (Fig. 3).

How does the net noise propagation to the target protein behave? The net propagation of noise from TF to target protein can be measured by 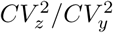. It is interesting to see that although 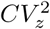 shows a distinct behavior, the propagation of noise shows enhancements for both the cases (Fig. 3(D)). However, it should be noted that noise propagation is significantly higher for the step-like dose response.

To better understand these results we decided to focus on the timescale of fluctuations of the free TF level in the presence of decoys. We computed the auto-correlation function of the free TF level using stochastic trajectories at steady-state as per

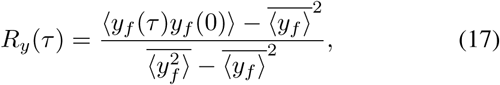

A slower decay of *R*_*y*_(*τ*) enhances noise propagation but a faster decay reduces noise propagation [25], [52]. In Fig. 4, we plot auto-correlation function for *N* = 0, 1000, and 3000 revealing that the decay of *R*_*y*_ becomes slower for larger decoy abundances, explaining the enhancement in the noise propagation seen in both the linear and step-like dose responses (Fig .3). Why is the noise propagation higher for the step-like dose response? Note that for a step-like dose response, noise propagation to the target protein only depends on the speed of fluctuations in *y*_*f*_ (*t*), i.e., how fast TF levels go below the threshold and bounce back. In contrast, for a linear dose response, noise propagation depends both on the speed of fluctuations and the magnitude of fluctuations in the free TF level. As the fee TF noise levels decrease with increasing *N* (Figs. 2 and 3), they buffer the noise propagation in the linear case, but not in the step-like nonlinear case.

**Fig. 4.**
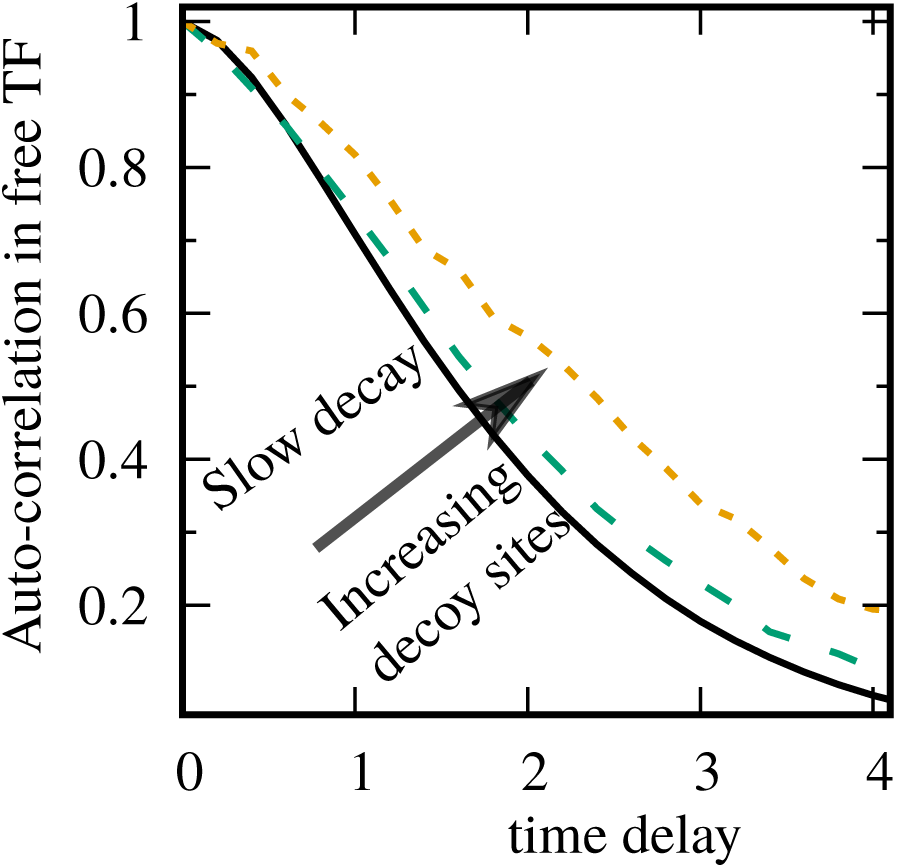
Decoy slow the speed of the fluctuations in the levels of the free TF. Auto-correlation function *R*_*y*_ for free TF as a function of time for *N* = 0, 1000, and 3000. The decay of *R*_*y*_ becomes slower for larger *N*. Parameter used: ⟨*B*_*x*_⟩ = 50, *k*_*x*_ = 1, *k*_*y*_ = 50, *γ*_*x*_ = *γ*_*y*_ = 1, and *k*_*d*_ = 1.

## IV. Conclusion

A significant portion of gene expression noise is extrinsic and can propagate downstream to target proteins with important consequences for cellular functioning. We have investigated the role of decoy binding sites on the propagation of extrinsic noise to downstream proteins. For this, we have formulated a stochastic model where a dynamical extrinsic disturbance modulates the synthesis rate of a TF, and free TF molecules can bind to genomic decoy binding sites that are present in a fixed number.

We have obtained analytical formula of noise in free TF counts in the limit of fast binding/unbinding, assuming small fluctuations around the mean. We have observed that free TF noise levels decrease as a function of the total number of decoy sites, and approach the Poisson limit for large decoy abundances (Fig. 2). Interestingly, the noise suppression ability of decoys is highest at intermediate timescales of disturbance fluctuations.

Finally, we have analyzed noise propagation to a target protein by stochastic simulations. Surprisingly, the noise in the target protein shows an enhancement with the addition of decoy sites when its activation by a TF occurs digitally beyond a threshold (Fig. 3). Thus, while increasing decoys decrease noise in the free TF level, it increases noise in the target protein. We argue that this increase in noise is explained by a slowing of fluctuations in the free TF level (Fig. 4), and hence the random switch between turning target proteins expression on and off occurs more slowly. A key focus of future work will be to develop theoretical tools to quantify noise propagation and systematically explore them for different dose-response shapes.

## ACKNOWLEDGMENT

This work is supported by the National Science Foundation Grant ECCS-1711548.

